# Multioviz: an interactive platform for *in silico* perturbation and interrogation of gene regulatory networks

**DOI:** 10.1101/2023.10.10.561790

**Authors:** Helen Xie, Lorin Crawford, Ashley Mae Conard

## Abstract

In this paper, we aim to build a tool that will help bridge the gap between high-dimensional computation and wet-lab experimentation by allowing users to interrogate genomic signatures at multiple molecular levels and identify best next actionable steps for downstream decision making. We introduce `Multioviz`: a publicly accessible `R` package and web application platform to easily perform *in silico* hypothesis testing of generated gene regulatory networks. We demonstrate the utility of `Multioviz` by conducting an end-to-end analysis in a statistical genetics application focused on measuring the effect of *in silico* perturbations of complex trait architecture. By using a real data set from the Wellcome Trust Centre for Human Genetics, we both recapitulate previous findings and propose hypotheses about the genes involved in the percentage of immune CD8+ cells found in heterogeneous stocks of mice. Source code for the `Multioviz R` package is available at https://github.com/lcrawlab/multio-viz and an interactive version of the platform is available at `multioviz.ccv.brown.edu`.

## Introduction

Phenotypic architecture is often driven by a collection of biological processes that occur through dynamic interactions across various molecular levels, including single nucleotide polymorphisms (SNPs), genes, and proteins [1]. Gene regulatory networks (GRNs) are directed graphs that effectively allow for the visualization of interactions between these components that constitute cellular pathways and signaling cascades [2]. Each node in a GRN represents a molecular variable such as a SNP or a gene, with each edge representing the interaction between two nodes. By simultaneously characterizing phenotypes at multiple genomic levels and modeling their interactions as a GRN, practitioners and data analysts can identify significant molecular variables for follow-up studies (e.g., through knockout experiments) [3].

Unfortunately, finding cost effective ways to investigate how a set of perturbations on a GRN will drive changes within a phenotype remains a challenge — especially as sequencing technologies continue to advance and, with this new depth, the space of potential biomarkers continues to grow. This motivates using *in silico* approaches to explore initial hypotheses and to identify actionable candidates for downstream tasks. To date, many computational methods have been developed for this purpose in high-dimensional multi-omics data settings [4, 5]. These tools often leverage variable importance measures, such as p-values and posterior inclusion probabilities (PIP), to infer GRNs [6, 7]. However, despite their usefulness, the software accompanying these algorithms usually require a fair amount of coding expertise to run them — thus, posing a challenge particularly for non-computational users [8]. Furthermore, the outputs of these methods are usually just lists of potential biomarker candidates which are not always easily amenable for determining the best next experimental action. To that end, there is a need for an accessible interactive platform that leverages statistical variable selection methods and subsequently enables non-computational researchers to efficiently test biological hypotheses *in silico*, prior to spending time and money in the wet-lab.

To meet this need in the field, we developed `Multioviz`: a web-based tool and `R` package for *in silico* exploration and assessment of GRNs. While many GRN platforms have been developed, a majority do not allow for perturbation analyses where a user is able to impose modifications onto a network (i.e., the addition or subtraction of a node or edge) and invoke a statistical reanalysis to learn how a phenotype might change with new sets of molecular interactions [9–13]. More notably, existing platforms that do indeed have the capability to incorporate perturbation analyses, often do not offer a user-friendly interactive environment for efficiently visualizing changes to GRNs [14, 15]. The key contribution of `Multioviz` is that it enables *in silico* perturbation experiments within an easy-to-use interface that includes the following three main features (Table 1). First, it allows users to couple summary statistics from a computational analysis (e.g., p-values or PIPs) along with a set of biological annotations (e.g., SNPs within the boundary of a gene) to visualize multi-level genomic relationships in the form of a GRN. Second, it allows users to perturb these learned networks and and investigate the associated ramifications on a phenotype of interest. Lastly, `Multioviz` integrates various variable selection methods to give users a wide choice of statistical approaches that they can use to generate relevant multi-level genomic signatures for their analyses.

**Table 1.**
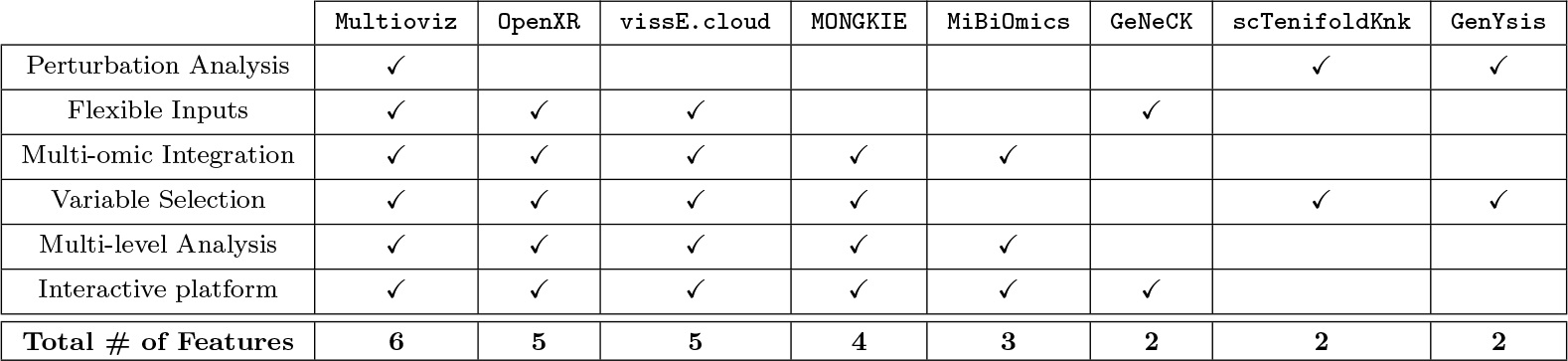
`Multioviz` combines the features of existing platforms to present a unified tool for gene regulatory network (GRN) based *in silico* hypothesis testing and perturbation analyses. Comparable platforms listed include: `OpenXR` [9], `vissE.cloud` [10], `MONGKIE` [11], `MiBiOmics` [12], `GeNeCK` [13], `scTenifoldKnk` [14], and `GenYsis` [15].

Overall, `Multioviz` provides an intuitive approach to *in silico* hypothesis testing, even for individuals with less computational and coding experience. Here, a user starts by inputting molecular data along with an associated phenotype to graphically visualize the relationships between significant variables. The user can then “knock out” a node in the GRN and rerun the statistical variable selection step to observe the effect of the perturbation. As a general illustration of our proposed platform, we will demonstrate how to perform perturbation analyses with the `Multioviz` online tool using “Biologically Annotated Neural Networks” (BANNs) which are a class of feedforward Bayesian machine learning models that integrate known biological relationships to perform association mapping on multiple molecular levels simultaneously [7]. The rest of the paper is organized as follows. In the next section, we describe the methodological and engineering details behind the three main features of `Multioviz`. Next, we demonstrate how to perform perturbation analyses using `Multioviz` with real quantitative traits assayed in a heterogeneous stock of mice from Wellcome Trust Centre for Human Genetics [16]. Finally, we close with a discussion and a look towards future research directions. We believe that the `Multioviz` platform and its application are a step towards providing practitioners the ability to perform true human-in-the-loop assessment of the biological processes driving complex phenotypes and diseases.

## Materials and Methods

The `Multioviz` platform allows the user to (i) intuitively visualize gene regulatory networks (GRNs) from multi-omics data (ii) perform *in silico* hypothesis testing through perturbing those GRNs and uncovering the effect the phenotypic architecture and (iii) allows these features to be leveraged with virtually any variable selection method. The general end-to-end workflow of `Multioviz` is intended to be intuitive and straightforward to all users regardless of coding experience (Figure 1). To begin, a user first inputs individual level data or summary statistics derived from a multi-omic data set (Figure 1a-b). In this paper, for the second step, we will demonstrate `Multioviz` using BANNs to perform variable selection on these input data. After statistically significant variables are identified, `Multioviz` outputs a GRN where the nodes correspond to genomic units (e.g., SNPs or genes) and the edges between nodes symbolize that there is some functional relationship that connects them. The BANNs method produces a posterior inclusion probability (PIP) for each molecular variable. These PIP scores lie on the unit interval and provide a prioritization score for each genomic variable in the data — with values closer to 1 indicating greater statistical significance [17]. In the `Multioviz` user-interface, these PIP values are displayed in different colors for nodes and edges, respectfully, which provides an interpretable view of important molecular variables. Insignificant SNPs are appear yellow, progressing to more red for those that are significant. Similarly, insignificant genes appear as light blue and then progress to dark blue for more significant genes (Figure 1c). In the third step of the `Multioviz` workflow, the user then has the flexibility to perturb any part of the GRN within the interface (e.g., by adding or removing variable nodes from the graph) to investigate *in silico* hypotheses (Figure 1d). The user can subsequently click a button to rerun the statistical analyses (e.g., BANNs in web application or another variable selection method in the `Multioviz R` package), and observe the newly visualized GRN (Figure 1e). The human-in-the-loop perturbation analyses provided by `Multioviz` will hopefully lead to better informed hypotheses to be tested and validated in the wet-lab for downstream tasks.In this section, we describe the GRN visualization, perturbation, and `R` package features in more detail.

**Figure 1.**
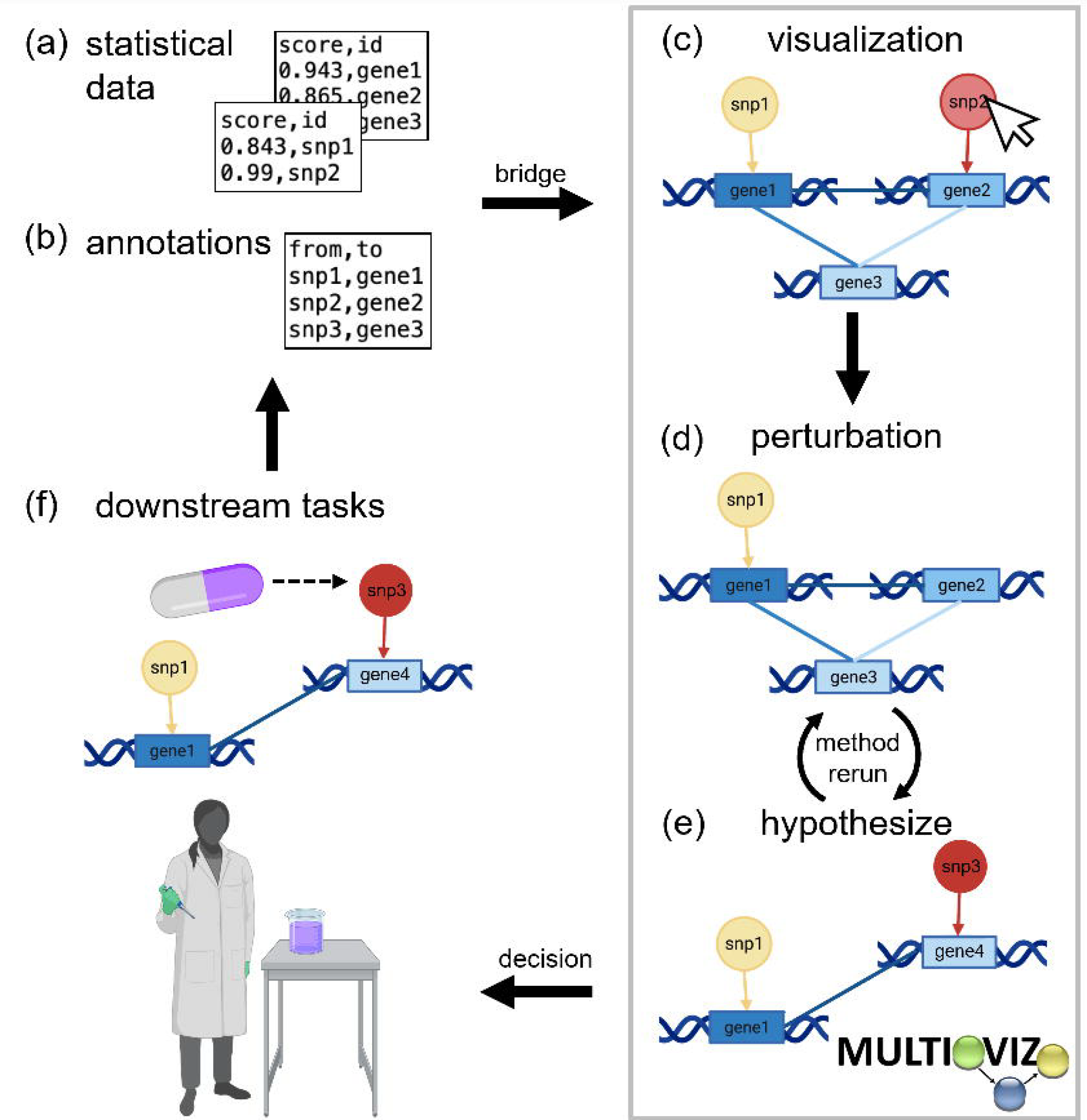
Schematic overview of running an end-to-end computational analysis with the Multioviz platform. **(a-b)** The user uploads their own individual-level data or summary statistics derived from an omics study. **(c)** Input data are visualized as gene regulatory networks (GRNs). Here, darker node colors denote greater statistical significance for a genomic variable. The mapping within and between molecular levels are given via edges which share the same color as the out-degree node. **(d)** `Multioviz` allows users to visualize and perturb GRNs from a prioritized list of significant molecular variables. **(e)** Through the perturbation feature, users can explore their generated GRN and delete nodes, then rerun statistical analyses to produce a new GRN. Subsequent rerunning of the variable selection method regenerates data to be visualized as an updated GRN. **(f)** Human-in-the-loop perturbation analyses provide better informed *in silico* hypotheses to be tested and validated in the wet-lab.

### Interpretable Visualization of Gene Regulatory Networks

The first step of the `Multioviz` workflow is to visualize molecular variables in the context of a GRN (Figure 2). The minimum required input is a file with two columns: (i) `id` which lists the molecular variables of interest and (ii) `score` which provides an associated summary statistic for each. `Multioviz` directly visualizes these data as a GRN. The shape of each node in the GRN corresponds to the molecular level (e.g., a SNP versus a gene) and the node color represents the importance of each variable (e.g., PIP or p-value) (Figure 2a). Overall, variables with greater degrees of significance are plotted with darker colors. For the purposes of demonstrating `Multioviz` functionality, we illustrate multio-omic data with SNPs and genes as the two molecular levels. `Multioviz` color-codes the first (SNPs) and second molecular levels (genes) as yellow to orange circles and light to dark blue rectangles, respectively. Because `Multioviz` leverages biological annotations that allow for the inference of biological hierarchies, there are directed edges between molecular levels. For example, given that SNPs can occur within the boundary of a gene and affect its function, we represent the interaction between the two levels as directed arrows when moving from the SNP to gene level [18]. Edges within the same molecular level are undirected, since we do not assume to have information on temporality. To summarize, for a given GRN, there are three types of edge connectivities: (1) no connectivity between the nodes belonging to one molecular level because those variables do not interact biologically, (2) sparse connectivity, or (3) complete connectivity. In Figure 2b, we show no connections on the SNP-level and complete connectivity between genes. For a given GRN, the user can then select a subset of molecular variables to perturb (i.e., add or delete) and rerun the method to identify which variables are significant in this new biological context (Figure 2c).

**Figure 2.**
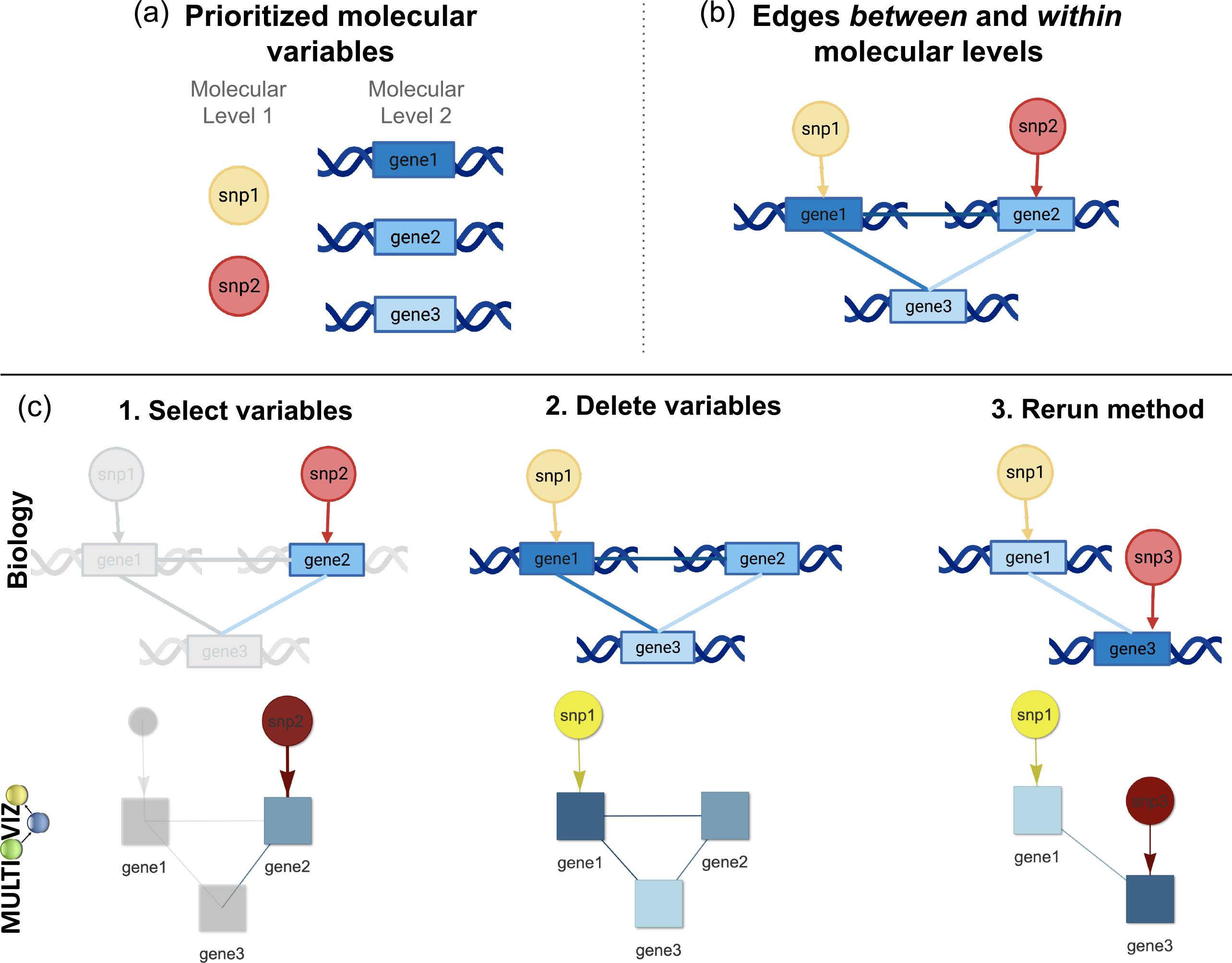
Components of gene regulatory networks (GRNs) and a schematic overview for performing perturbation analyses in Multioviz. **(a)** Visualization of the components making up a GRN. Here, variables (e.g., SNPs and genes) are represented as nodes. The shape of these nodes are different depending on the molecular level that they represent and the color scheme describes the significance level of each variable according to a statistical model (e.g., p-value or PIP). Insignificant SNPs are more yellow and more statistically important SNPs are depicted in red. Similarly, insignificant genes are light blue, while significant genes are dark blue. **(b)** Directed edges are used to map nodes between molecular levels (e.g., SNPs reside in the boundaries of genes) [18]. Since we do not assume to have access to temporal information, interactions between variables on the same molecular level are represented by undirected edges. There are three classes of edges between and within molecular levels: no connectivity, sparse connectivity, and complete connectivity. Here, the first molecular level has no connectivity since there are no direct interactions between SNPs. However, the second molecular level has complete connectivity because there is an interaction between all genes. The nodes and edges together form a visual representation of a GRN. **(c)** To emulate perturbation analyses, `Multioviz` allows users to select molecular variables (i.e., nodes), delete them, and rerun the statistical analysis to generate a new GRN. To perform this type of analysis within in the `Multioviz` interface, users simply highlight the variable of interest by (1) clicking on the node and selecting “Edit”, (2) clicking on “Delete selected”, and then (3) clicking “Rerun” under the “Perturb” left drop-down menu. An overview of the `Multioviz` interface can be found in Figure 3.

When working directly in the `Multioviz` interface, users can click the “Visualization” left-hand side drop-down to upload their own file of statistical importance scores for variables up to two molecular levels (Figure 3a). This file should be a two-dimensional matrix with the column names labeled as “score” and “id”, respectively. Once these inputs are uploaded, the user can click “Run” to construct and view a corresponding GRN (Figure 3b). Note that licking on a specific node will highlight the variable itself along with its connected neighbors. If desired, the user can also upload their own biological annotations to define *a priori* relationships and generate sparse edges between genomic levels (e.g., SNPs-to-genes or genes-to-pathways). To further explore the output of the GRN, `Multioviz` offers the user the flexibility to set importance thresholds for each molecular level and filter out variables with low significance (Figure 3c). This can be particularly important when generating GRNs from large data sets with many variables. Finally, users can manually modify the GRN after it is generated to create layouts that are most digestible (Figure 3d).

**Figure 3.**
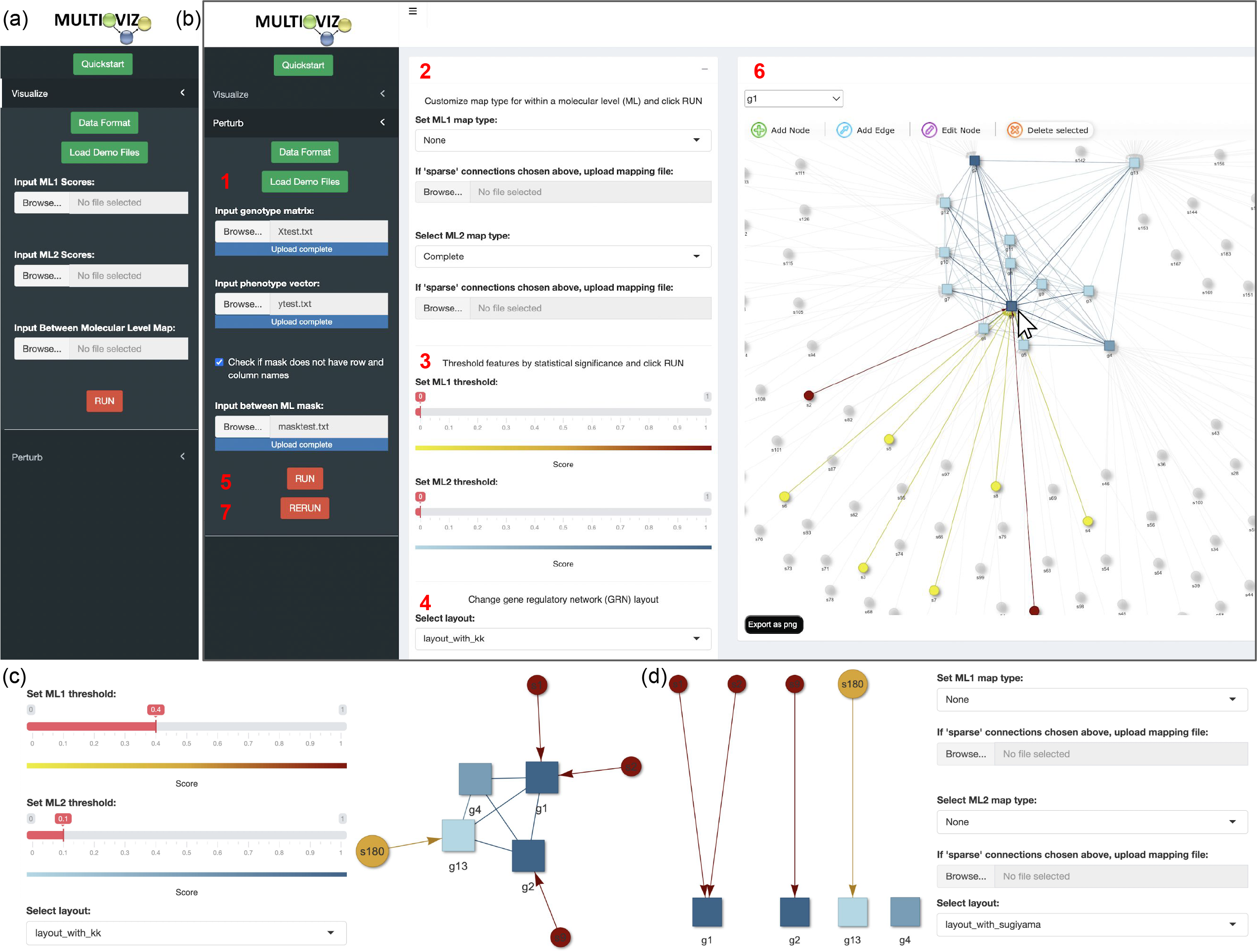
Overview of the Multioviz online tool user interface. **(a)** Users can select the visualization drop down to upload pre-generated variable ranks and annotation maps between molecular variables. **(b)** Under the perturbation drop down, the user can follow the outlined steps numbered in red. In step (1), the user will upload the required genotype matrix, phenotype vector, and biological annotation map (often given as a binary matrix where a “1” means that a variable belongs to a given group). In steps (2) through (4), the user can customize the type of method being used for association mapping, the threshold used to determine variable significance, and the layout for their gene regulatory network. In step (5) run BANNs (or some other variable selection approach) and subsequently generate the GRN. In step (6), the user can perform a perturbation analysis where they can select and delete variables of interest. Finally in step (7), the user will rerun the method to test *in silico* hypotheses. **(c)** `Multioviz` allows the user to adjust significance thresholds for each molecular level. **(d)** The user is able to specify the degree of mapping within each molecular level, thereby changing and/or modifying the GRN layout.

### Facilitated Perturbation Analyses for *in silico* Experimentation

The flexible functionality of `Multioviz` allows for the *in silico* testing of hypotheses where the nodes and edges of a learned GRN can be perturbed to observe the influence of different molecular variables onto a phenotype of interest. These changes are run as altered inputs in a variable selection model that runs in the background of the software which then generates a new set of significant molecular variables that are then visualized as another interactive GRN (see Figure 3). In the `Multioviz` web application, the statistical method that is implemented is BANNs [7]. Like many linear and nonlinear models, BANNs requires a genotype matrix and a phenotype vector as input. Consider a biological study with *N* observations (e.g., the number of individuals, cells, tissues) that have been phenotyped for some response **y** = (*y*_1_, …, *y*_*N*_). Assume that the *i*-th sample has been genotyped, sequenced, or profiled for *J* variables (*x*_*i*1_, …, *x*_*iJ*_) (e.g., gene expression, single nucleotide polymorphisms, proteomics). Collectively, all variables across all samples can be collected in an *N* × *J* matrix **X**.In the `Multioviz` interface, users can click the “Perturb” drop down menu and upload their data *D* = {**X, y**} along with a set of biological annotations encoded as a *J* × *G* binary mask matrix **M** where *G* denotes the number of groups on the second molecular level. In this case, we have *J* SNPs that are grouped into *G* genes. To run BANNs (or a similar variable selection approach) and generate a GRN, the user should click “Run” once their data are uploaded. Once the user has a clear understanding of the GRN, *in silico* hypothesis testing can carried out by clicking on a variable of interest, selecting “Edit” to delete the node and its connected edges, and then clicking “Rerun” to rerun BANNs and generate new sets of interactive GRN. Overall, this human-in-the-loop process facilitates the efficient testing of any number of hypotheses.

### Flexible Integration of Statistical and Machine Learning Methods

Part of the contribution of `Multioviz` is that it is also available as a standalone `R` package. This enables users with coding experience to have more control over the statistical methodologies that run in the background of the software. This is important when there are unique theoretical considerations that need to be made for different types of omic data before performing perturbation experiments. Regardless of the method used, we recommend that the model have the ability to variable selection or regularization to ensure that the resulting GRN is reasonably sized (i.e., reducing an initial high-dimensional set of variables to a small number worthy of follow-up). Implementing `Multioviz` within a developer script only requires two inputs once the package is installed: (i) a file of molecular variables and their associated scores, and (ii) a set of biological annotations.

## Results

To demonstrate the utility of `Multioviz`, we apply the software to real genetic data from a heterogeneous stock of mice collected by the Wellcome Trust Centre of Human Genetics (http://mtweb.cs.ucl.ac.uk/mus/www/mouse/index.shtml) [16]. The genotypes from this study were downloaded directly using the `BGLR R` package [20]. This study contains *N* = 1,814 heterogeneous stock of mice from 85 families (all descending from eight inbred progenitor strains) and 131 quantitative traits that are classified into 6 broad categories including behavior, diabetes, asthma, immunology, haematology, and biochemistry.

Phenotypic measurements for these mice can be found freely available online to download (details can be found at http://mtweb.cs.ucl.ac.uk/mus/www/mouse/HS/index.shtml). In this study, we focus on modeling the percentage of CD8+ cells in these mice as our **y** vector. For preprocessing, we corrected this trait for sex, age, body weight, season, and year [16]. The **X** matrix that we input into `Multioviz` contains single nucleotide polymorphisms (SNPs) as variable, each of which are encoded as {0, 1, 2} copies of a reference allele at each locus. For mice with missing genotypes, we imputed values by the mean genotype of that SNP in their corresponding family. Only polymorphic SNPs with minor allele frequency above 5% were kept for the analyses. This left a total of *J* = 10,227 SNPs that were available for all mice. Lastly, to create biological annotation file **M**, we used the Mouse Genome Informatics database (http://www.informatics.jax.org) [21] to map SNPs to the closest neighboring gene(s). Unannotated SNPs located within the same genomic region were labeled as being within the “intergenic region” between two genes. Altogether, a total of *G* = 2,616 annotations were analyzed.

We input these files into `Multioviz` where we assumed that significant SNPs and genes would produce PIPs greater than or equal to 0.5 — this is also known as the median probability model threshold in Bayesian statistics [19]. When viewing the corresponding GRN produced by the software, this resulted in 15 associated SNPs variables and 19 enriched genes (Figure 4). Notably, we observed the SNP *CEL-17_31069801* and gene *hlb156* on chromosome 17 as both being significant (PIPs = 1). As corroborating evidence, the genomic region where these molecular variables reside has been reported to contain highly significant SNPs that contribute to non-additive variation for CD8+ T-cells [16]. To investigate this region further, we perturbed the GRN in `Multioviz` by deleting *CEL-17_31069801* and observed the emergence of *CEL-17_31214920* as being important which also maps to the *hlb156* gene (PIP = 1). The two new gene-level variables that also became enriched upon perturbation are both associated with CD8+ T-cell differentiation: *Anapc1* (PIP = 0.726) and *Pard3* (PIP = 0.998). *Anapc1* functions in the metaphase-to-anaphase transition in the cell cycle and has been associated with poor prognosis in T-cell acute lymphoblastic leukemia [22]. *Pard3* directs polarized cell growth and asymmetric cell division [23]. The asymmetric division of T-cells has been uncovered as a potential means by which effector and memory T cells are differentiated during immune responses [24]. Overall, we show here that `Multioviz` has the potential to enable users to generate new testable hypotheses *in silico* through its perturbation framework. These results suggest a honed set of molecular variables to explore in investigating mechanisms underlying the percentage of CD8+ T cells in heterogeneous mice.

**Figure 4.**
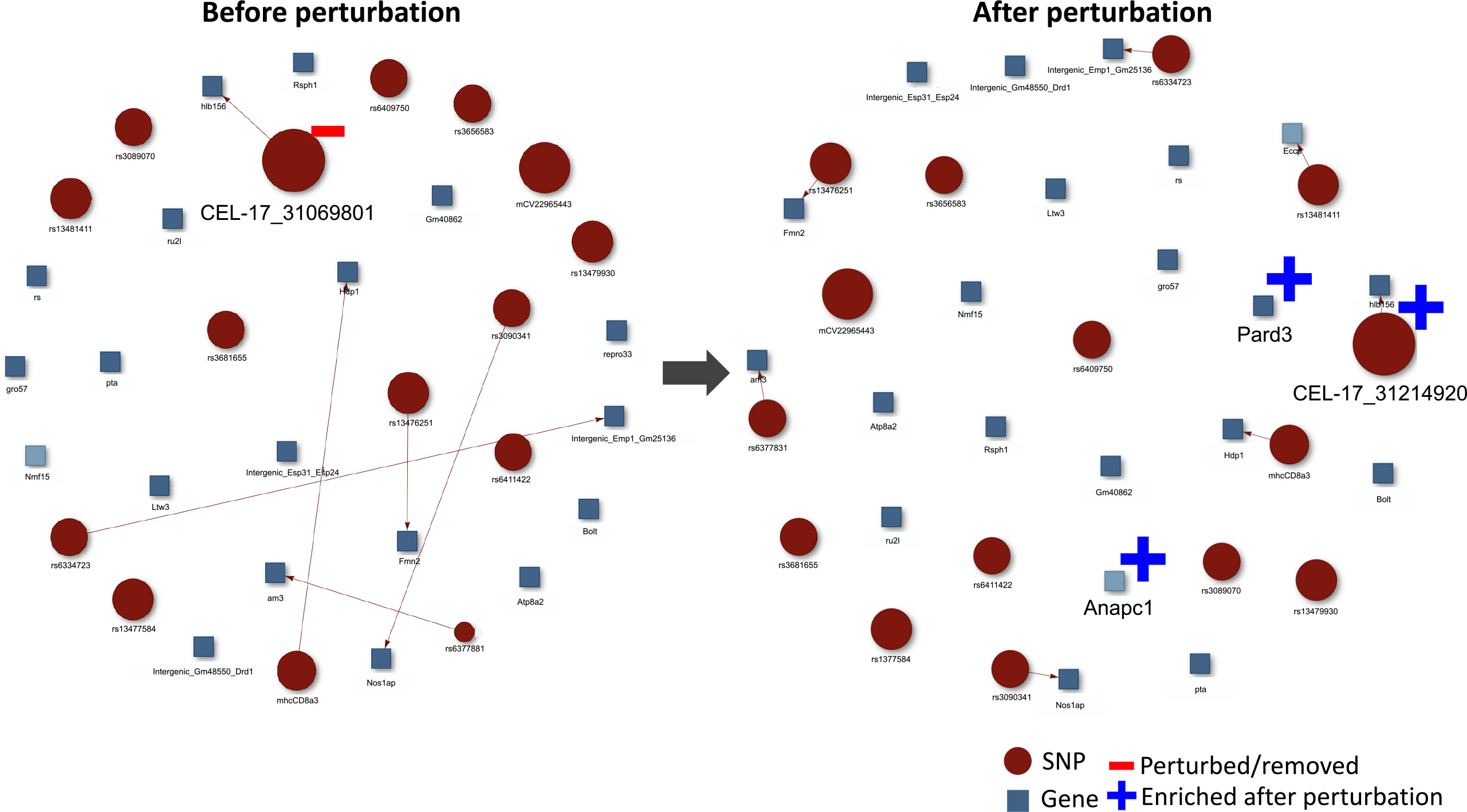
Demonstration of an *in silico* analysis using Multioviz on a heterogeneous stock of mice data set from the Wellcome Trust Centre of Human Genetics. We applied `Multioviz` to visualize a GRN with associated SNPs and enriched genes driving the architecture of CD8+ cell percentage. To generate this GRN, we set the following parameters in the `Multioviz` software: (i) `Molecular Level 1 (ML1) Map Type = None`; (ii) `Molecular Level 2 (ML2) Map Type = None`; (iii) `ML1 Threshold = 0.5` which follows the median probability model [19]; (iv) `ML2 Threshold = 0.5`; and `GRN Layout = “layout_with_kk”`. In this figure, SNP-level variables in ML1 (red circles) map to gene-level variables in ML2 (blue squares). Upon deleting the *CEL-17_31069801* SNP and rerunning `Multioviz`, we observe a new association with the SNP *CEL-17_31214920* and a new enrichment of the genes *Anapc1* and *Pard3*. These are depicted in the perturbed GRN on the right. Note that both the deleted and newly enriched SNPs map to the *hlb156* gene.

## Discussion

`Multioviz` is an interactive platform for *in silico* hypothesis testing with GRNs. Both the web platform and the `R` package allow users to easily explore interactions between variables in omics data sets through clear visualizations and by enabling them to perform perturbation analyses. It is well known how valuable it can be to perform *in silico* knock-out or knock-down experiments to determine the best next actionable steps, prior to performing follow-up *in-vivo* and *in-vitro* experiments [25]. The `Multioviz` platform is in service of this goal. Our real data results with the heterogeneous stock of mice data set from the Wellcome Trust Centre for Human Genetics serve as an illustrative example of how `Multioviz` can be used to identify a small set of candidate molecular variables that could be implicated in CD8+ T-cell differentiation. As part of future work, we want to extend `Multioviz` to integrate a wider array of mathematical and machine learning models, as well as allow for integrating more that just two molecular levels for analysis.

Overall, we envision tools like `Multioviz` being used for applications such as early drug development where the goal is often not only to identify potential druggable targets for disease pathways but also test the effects of drugs *in silico* prior to moving them into a biological system. Further, tools including `Multioviz` could serve as a powerful means through which clinicians can generate and explore GRNs on a patient level and, as such, prescribe treatments and dosages tailored to each patient. The `Multioviz` platform is freely available, thereby providing researchers with an accessible way to analyze punitive molecular mechanisms underlying various traits across a wide array of biological levels.

## Software Availability

`Multioviz` is currently available as both a free web application and an `R` package. The web platform is hosted through the Center for Computation and Visualization at Brown University and can accessed at `multioviz.ccv.brown.edu`. The `R` and other source code are also freely available at https://github.com/lcrawlab/multio-viz. This GitHub repository also provides additional robust details on the platform and extensive tutorials on how to run analyses in the README.

## Acknowledgements

This research was conducted using computational resources and services at the Center for Computation and Visualization (CCV), Brown University. We would like to thank Brown’s Computational Biology Core for hosting `Multioviz` online and making it free for users. Components of Figure 1 (f) were created with BioRender.com.

